# In silico investigation of cytochrome bc1 molecular inhibition mechanism against *Trypanosoma cruzi*

**DOI:** 10.1101/2022.06.01.494283

**Authors:** Stefano Muscat, Gianvito Grasso, Leonardo Scapozza, Andrea Danani

**Author notes:** **Corresponding Author** Andrea Danani - Istituto Dalle Molle di studi sull'Intelligenza Artificiale (IDSIA), Scuola universitaria professionale della Svizzera italiana (SUPSI), Università della Svizzera italiana (USI), Lugano, Switzerland.

## Abstract

Chagas’ disease is a neglected tropical disease caused by the kinetoplastid protozoan *Trypanosoma cruzi*. The only therapies are the nitroheterocyclic chemicals nifurtimox and benznidazole that cause various adverse effects. The need to create safe and effective medications to improve medical care remains critical. The lack of verified *T. cruzi* therapeutic targets hinders medication research for Chagas’ disease. In this respect, cytochrome bc1 has been identified as a promising therapeutic target candidate for antibacterial medicines of medical and agricultural interest. Cytochrome bc1 belongs to the mitochondrial electron transport chain and transfers electrons from ubiquinol to cytochrome c1 by the action of two catalytic sites named Qi and Qo. The two binding sites are highly selective, and specific inhibitors exist for each site. Recent studies identified the Qi site of the cytochrome bc1 as a promising drug target against *T. cruzi*. However, a lack of knowledge of the drug mechanism of action unfortunately hinders the development of new therapies. In this context, knowing the cause of binding site selectivity and the mechanism of action of inhibitors and substrates is crucial for drug discovery and optimization processes. In this paper, we provide a detailed computational investigation of the Qi site of *T. cruzi* cytochrome b to shed light on the molecular mechanism of action of known inhibitors and substrates. Our study emphasizes the action of inhibitors at the Qi site on a highly unstructured portion of cytochrome b that could be related to the biological function of the electron transport chain complex.

## Introduction

The protozoan parasite *Trypanosoma cruzi* causes the Chagas’ disease, a neglected tropical disease that is endemic in South and Central America. Chagas’ disease has also spread through migration across North America, Europe, and the Western Pacific region, infecting about 8 million people worldwide and causing 10·000 deaths annually(1,2). The treatment relies on the nitroheterocyclic compounds nifurtimox and benznidazole, effective in the acute phase but less so in the chronic phase. Many treatments for Chagas’ disease cause severe toxic effects and encounter drug resistance mechanisms(3,4). New effective therapies in advanced clinical development are scarce, and the need to develop safe and effective drugs to improve medical treatment remains urgent.

Drug discovery for the treatment of Chagas’ Disease is hampered by the scarcity of validated *T. cruzi* drug targets. In this context, cytochrome bc1 has been identified as a drug target for antimicrobial agents of medical and agricultural interest(5,6) and as a promiscuous drug target candidate for *T. cruzi*(7). The cytochrome bc1, also called cytochrome c oxidoreductase or complex III, is part of the electron transport chain (ETC) and catalyzes the transfer of electrons from ubiquinol to cytochrome c and couples electron transport and proton translocation across the inner mitochondrial membrane through the Q cycle(8–10). In detail, cytochrome bc1 is a multi-subunit dimeric protein complex consisting of cytochrome b with two b-type hemes, cytochrome c1 with a c-type heme, and the Rieske iron sulfur protein subunit. Cytochrome bc1 has two binding sites called Qo and Qi, whose substrates are ubiquinol (QH2) and ubiquinone (Q), respectively. The Q cycle consists of two half-cycles in which a molecule of ubiquinol binds consecutively in each half-cycle to the Qo site and is oxidized, giving up an electron to Rieske iron-sulfur (FeS) protein and an electron to low potential b heme (bL-heme). Consequentially, two protons are transferred into the intermembrane space. In turn, the electron arriving on the bL-heme group is used to reduce the high potential b heme (bH-heme). At the same time, a molecule of ubiquinone binds to the Qi site and is reduced by the bH-heme to form a semiquinone radical. In the next half-cycle, another electron reduces again the semiquinone radical to form ubiquinol, importing others two protons from the mitochondrial matrix.

Inhibition of the activity of the bc1 complex blocks the generation of ATP (11) and several inhibitors have been investigated in the year, specific for each binding site. Among the known inhibitors for the Qo site, myzothiazol and strobilurin A(12) block the reduction and oxidation pathway of ubiquinol on the iron-sulfur protein binding in the bL-heme proximal and bL-heme distal domain, respectively (13–16). In recent years, efforts have been concentrated to develop new generation of inhibitors that target the Qi site. In detail, several studies highlighted the antimycin A(17) action to block the respiratory activity targeting the Qi site. Furthermore, GNF7686(18) is one the most promising Qi site inhibitors, also effective against *Leishmania donovani* and *T. cruzi* (7).

The development of new drugs is often hindered by a lack of information about the mechanism of action. Furthermore, the recent failure of posaconazole, an established CYP51 inhibitor, in phase II clinical trials for Chagas’s Disease highlights the need to better understand the mechanisms of action of drug candidates(19,20). In this context, understanding the mechanisms of binding site selectivity and the mechanism of action of inhibitors and substrates is essential for drug discovery and drug optimization processes. The study of molecular action faces experimental technical limitations due to the complexity and metastable nature of the reaction complex. Computational modelling has been widely used to complement the experimental information of the mechanism at the atomistic level(21,22). Here, we used a comprehensive set of computational tools including homology modelling, molecular docking, and molecular dynamics simulations to investigate the mechanism of action of certain inhibitors and substrates specific of cytochrome bc1 with a focus on the Qi site. Specifically, we studied and evaluated the conformational changes that cytochrome b of *T. cruzi* undergoes when the Qi site is occupied by Qi inhibitors (antimycin A and GNF7686), Qo inhibitors (myxothiazol and strobilurin A), and substrates (ubiquinol and ubiquinone), comparing them to the apo-form.

## Materials and Methods

### Homology Modelling

The 3D structure of *T. cruzi* cytochrome b was not available in the literature and homology modelling techniques were employed to address this point. First, the Cytochrome b genes of *T. cruzi* were extracted from Silvio X10-1 genome (v39, tritrypDB) with maxi-circle (FJ203996.1, NCBI)(23). In order to obtain the amino acid sequence of *T. cruzi* cytochrome b, the RNA editing within the coding region before translation was necessary(24). The *T*.*cruzi* cytochrome b homology model was built using the crystallographic structure of the avian *Gallus gallus* (PDB ID: 1BCC)(25) selecting the chain C as a template as previously done in literature(7). The sequence alignments were performed using the T-Coffee algorithm(26) (see Figure S1). The two amino acids sequences exhibit the 45% of sequence identity of the Qi active site(7) and restraint-based homology modelling was performed using MODELLER software(27). Figure S2 shows a superposition of the avian Gallus model template of cytochrome b (PDB ID 1BCC – chain C) and the final *T. cruzi* homology model built, highlighting a good match of the two structures. The quality of the obtained structural model was evaluated by PROCHECK(28) observing the phi-psi angle distribution by a Ramachandran plot and more than 98% of the residues were in the most favored conformation regions, while less than 2% of residues were in disallowed regions (Figure S3).

### Molecular Docking

Molecular docking was carried out by PLANTS software(29). The ligands and the receptor were pre-processed by SPORES software(30). The PLANTS_CHEMPLP_ scoring function was used in combination with search speed 1, 20 aco ants, and 25 aco evaporation factor (29). The Qi binding center was set at the center of mass of antimycin A of the reference PDB 3H1I choosing a binding radius of 12 Å. The docking procedure was repeated 100 times selecting the pose with a higher affinity of each run. The best 100 poses were clustered based on the RMSD of the heavy ligand atoms. The final pose was the centroid with higher affinity. The robustness of the docking workflow was evaluated on the antimycin A pose (PDB ID 3H1I), obtaining RMSD below 2 Å (Figure S4). The docking procedure was repeated to evaluate three ligand classes bounded in the Qi site: Qi inhibitors (antimycin A and GNF7686), Qo inhibitors (myxothiazol and strobilurin A), and substrates (ubiquinol and ubiquinone). All molecular systems were compared to the cytochrome b apo-form.

### Molecular Dynamics Simulations

The cytochrome b – ligand complexes were inserted in a lipid bilayer of about 160 molecules of 1-palmitoyl-2-oleoyl-sn-glycero-3-phosphocholine (POPC) using CHARMM-Builder with 3 nm of water thickness and 150 mM NaCl to neutralize the net charge system. Each resulting molecular system was minimized by 1000 steps using the steep descent algorithm. Two 400 ps position restrained simulations with decreasing force constants to all heavy atoms (2000 and 1000 kJ mol-1·nm-2) were performed under NVT ensemble at 300 K using v-rescale thermostat (τ_T_ = 1.0 ps; time step = 1 fs)(31). Then, three position-restrained simulations of 500 ps (force constants of 1000, 400, 200 kJ mol^-1^·nm^-2^) were performed using Berendsen semi-isotropic pressure coupling scheme (time step = 2 fs; pressure time constant τP = 5.0 ps)(32) at a reference pressure of 1 atm. Finally, 100 ns of MD production was carried out using the V-rescale thermostat (T = 300 K; τ_T_ = 1.0 ps), together with the semi-isotropic Parrinello-Rahman barostat (P = 1 atm, τ_P_ = 2.0 ps)(33). The simulation time step was set to 2 fs in conjunction with the LINCS algorithm (34). The particle mesh Ewald (PME) method (35) was used to calculate electrostatic interactions, with a real-space cut-off of 1.2 nm and employing periodic boundary conditions. The van der Waals interactions were calculated by applying a cut-off distance of 1.2 nm and switching the potential from 1.0 nm. The topology parameters of the bH-heme molecules were taken from the literature(36), assigning the reduced state. The ligands forcefield topology was taken from the CHARMM General Force Field(37). Protein, phospholipids, water, and ions topology were parametrized by the CHARMM36m force-field(38). All MD simulations were carried out using GROMACS 2020.4 software package(39). The Visual Molecular Dynamics (VMD) software was used to monitor all simulation trajectories(40). ProLIF (41) package was used for ligand-receptor interaction analysis.

In order to increase the statistical power, three replicas of each molecular system have been carried out. An overview of the molecular systems investigated is reported in the Supporting Information (Table S1). The system analysis was carried out on the last 20 ns of each molecular system.

## Results

An extensive set of computational tools was used to investigate the structural conformational changes of the cytochrome b of *T. cruzi* evaluating three ligand classes bound in the Qi site: Qi inhibitors (antimycin A and GNF7686), Qo inhibitors (myxothiazol and strobilurin A), and substrates (ubiquinol and ubiquinone) (Figure 1). All molecular systems were compared to the cytochrome b apo-form.

**Figure 1.**
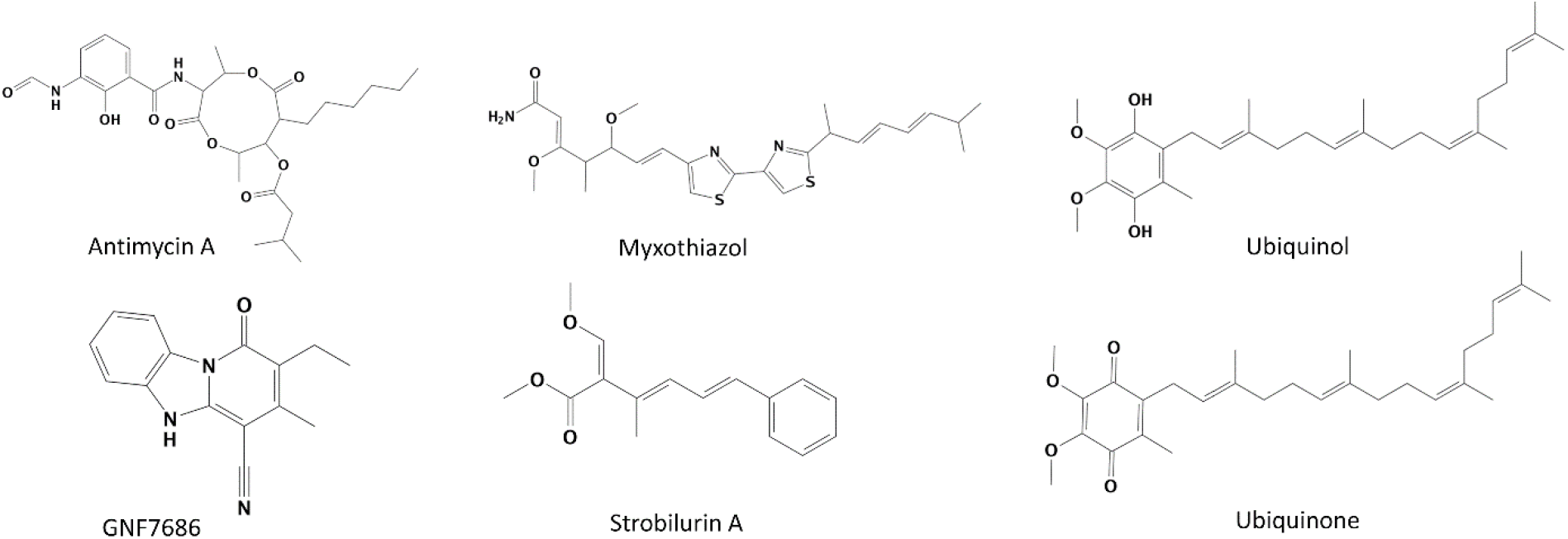
Chemical structures of Qi inhibitors (left), Qo inhibitors (middle), and substrates (right).

Molecular docking was used to predict the starting pose of each ligand at the Qi site. Figure 2 shows the predicted binding pose of each ligand. All the ligands are characterized by a hydrophobic tail distal to the bH-heme molecule. Regarding antimycin A, the planar benzene ring interacts with Leu197 and the middle carbonyl group with Gly30 and Phe33. GNF7686 occupies a smaller volume than the other ligands and is coordinated with Phe222. Myxothiazol exposes the amine group to the residues Leu197 and Leu25 and establishes interactions between thiazole rings and the residues Gly30 and Phe33. Strobilurin A interacts mainly with Leu25 by oxygen atoms and with Phe33 by the benzene group. Ubiquinol and ubiquinone have similar binding modes due to their similar chemical nature, interacting mainly with Leu197 and Leu200.

**Figure 2.**
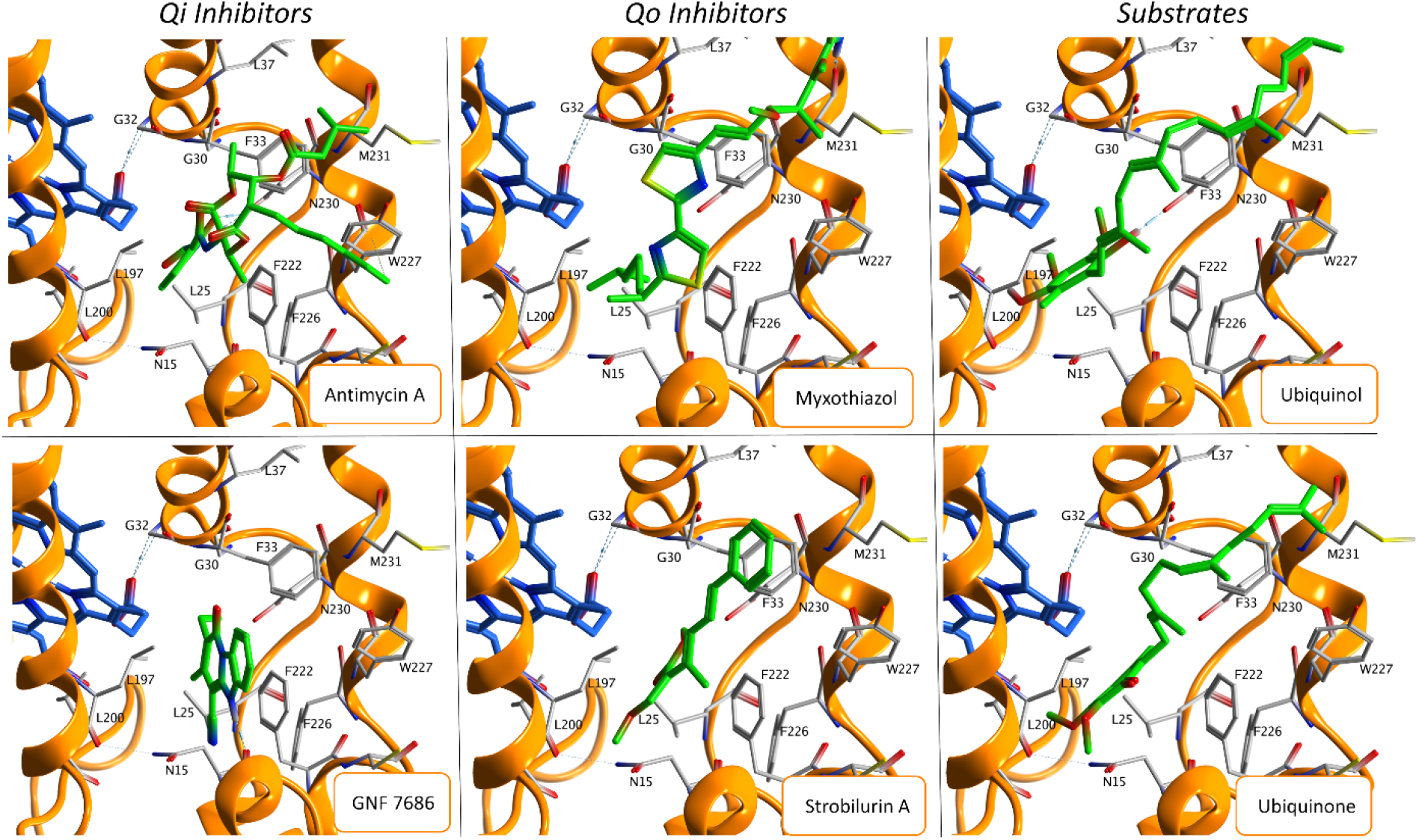
Best ligand centroids in terms of PLANTS scoring function of each ligand, divided in qi inhibitors, qo inhibitors and substrates. Each ligand is shown in green, the bH-heme molecule in blue, the cytochrome b in orange cartoon, and the main residues of the Qi pocket receptor in gray.

Each molecular docking configuration was used as starting point for MD simulations. To characterize the effect of ligand-cytochrome b interactions on protein conformational dynamics and the cytochrome b dynamics itself, three independent MD replicas have been performed for each molecular system. During the MD simulations, the cytochrome b’s Qi pocket undergoes deep conformational changes, and each ligand interacted with the Qi site in a different way. Figure 3 shows the bond probability between each ligand and the amino acids residues of the Qi pocket in the residues range 10 to 44 and 190 to 235. Overall, each ligand interacts with the residues of the Qi site mainly via hydrophobic interactions (blue lines, Figure 3). Antimycin A, myxothiazol, and the natural substrates can interact with a greater number of residues than other ligands, suggesting a greater compactness of the binding site. All ligands, except strobilurin A, form hydrogen bonds (yellow and green lines, Figure 3), mainly with residue Leu16 while antimycin A with Asp15. In addition, the Qi and Qo inhibitors also form Pi-stacking interactions (red lines, Figure 3) with phenylalanine residues: antimycin A with His202 and Phe222, myxothiazol and strobilurin A with Phe33 and GNF7686 with Phe194.

**Figure 3.**
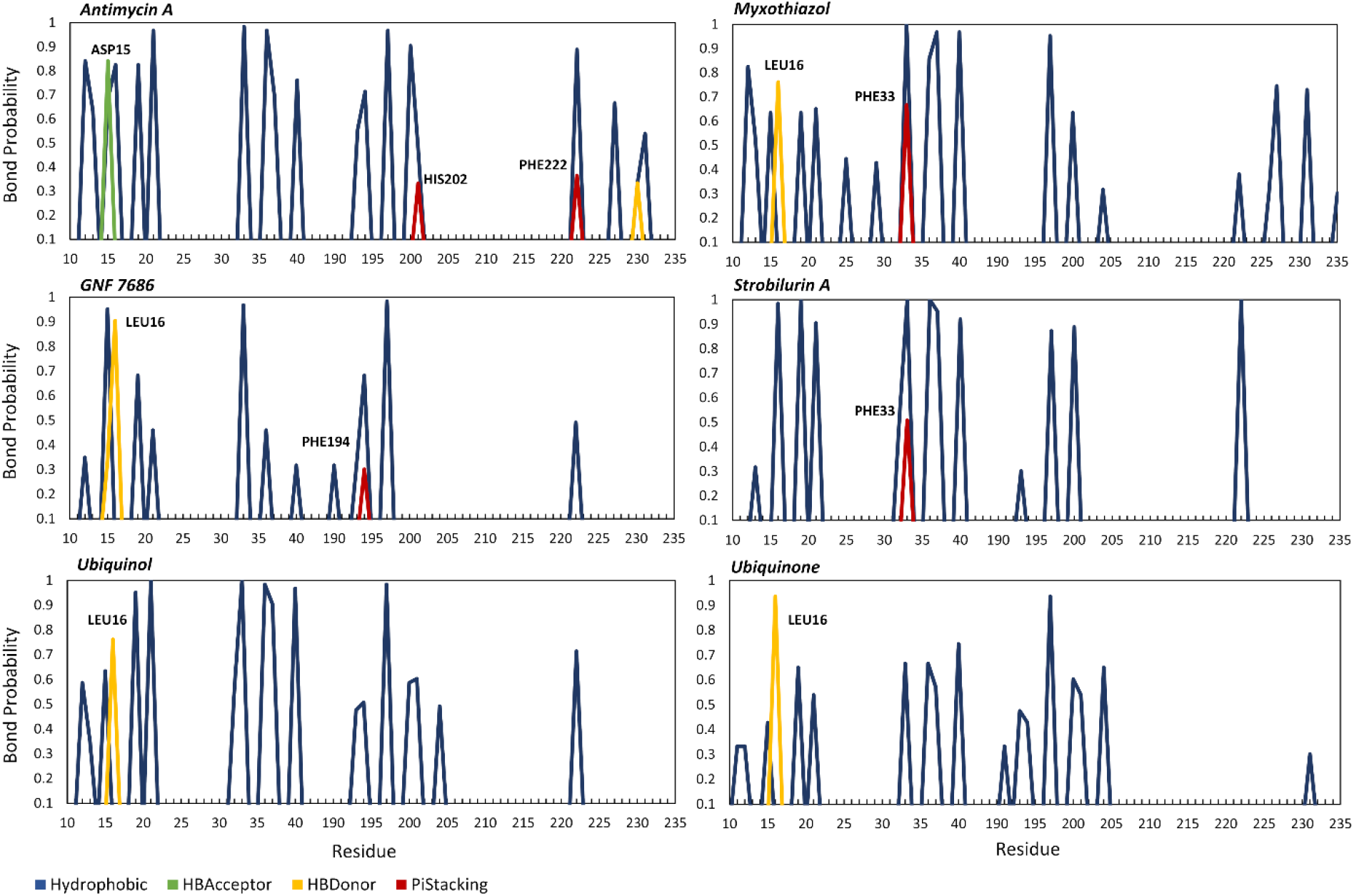
Bond probabilities between each ligand investigated and Qi receptor residues in the ranges F10 to I44 and I190 to F235.

Each class of ligands generates a conformational response of the Qi pocket that affects the conformation of a specific area of the protein. Figure 4 shows the root mean square fluctuation (RMSF) evaluated on the Cα atoms of the cytochrome b and has been calculated for each system, for each replica, and then reported as mean and standard deviation. The RMSF investigation highlights the largest change in dynamics of the cytochrome b receptor bounded with three classes of ligands and compared with the apo-form. Overall, approximately 92% of the residues of ligand-receptor complexes have a fluctuation comparable to the apo-form. The most important difference in term of RMSF is located within the protein domain L200-D230. When Qi inhibitors and the substrate ubiquinone are located at the binding site, the receptor residues exhibit fluctuation amplitudes similar to those of the apo-form. In contrast, when Qo site-specific inhibitors or the ubiquinol substrate are present at the Qi site, the residues range L200-D230 under investigation became more flexible, achieving RMSFs of 0.37 nm and 0.77 nm respectively. The analysis highlights a marked decrease of conformational fluctuations between the residues L200 and D230, when specific inhibitors of the Qi site and ubiquinone are bounded compared to the Qo inhibitors class and ubiquinol. The conformational changes to which the receptor is subjected may be related to the biological functioning of cytochrome bc1.

**Figure 4.**
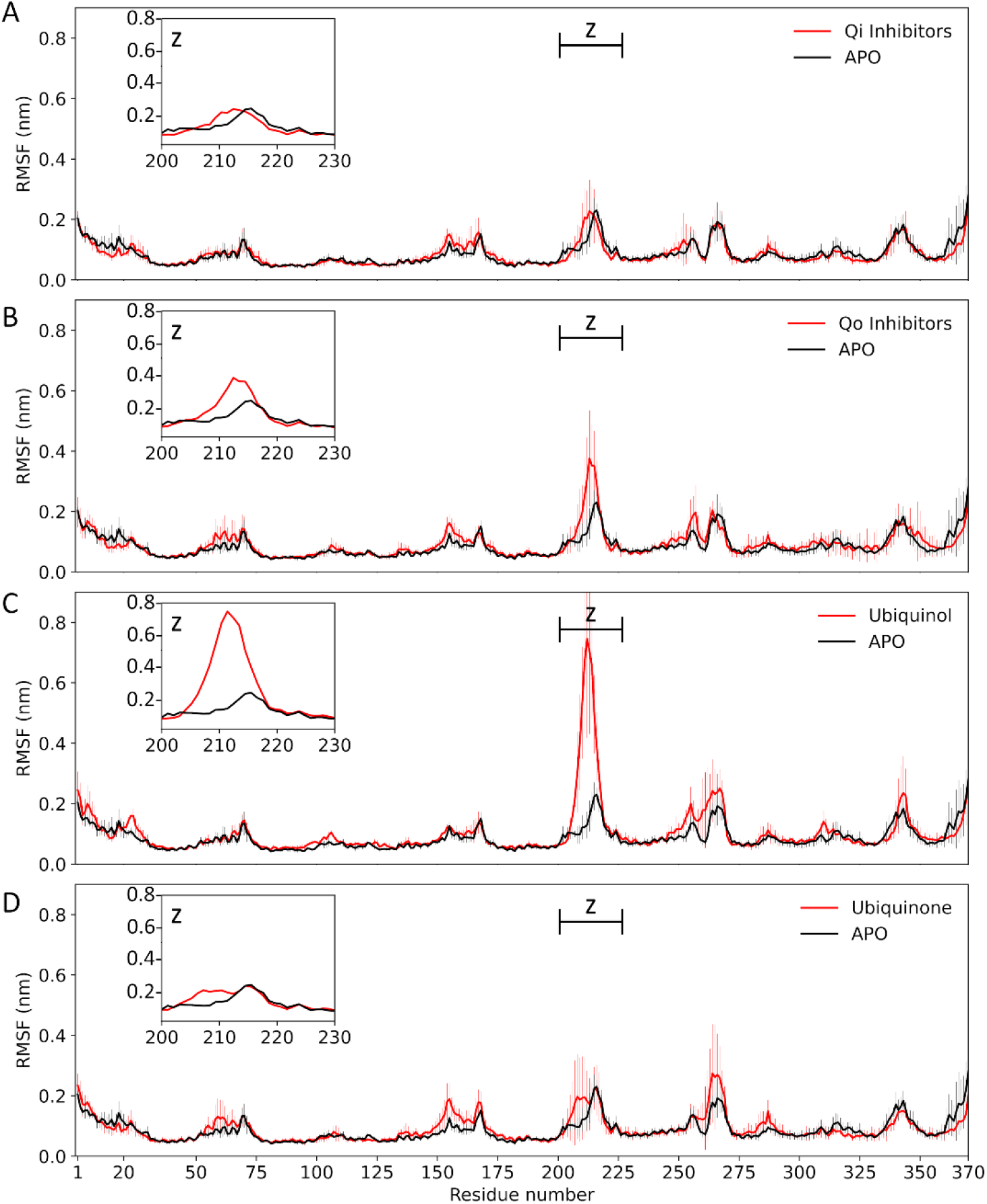
Root mean square fluctuation of cytochrome b backbone in presence of Qi inhibitors (panel A), Qo inhibitors (panel B), and two substrates (Ubiquinol in panel C and Ubiquinone in panel D) compared to the apo-form. The upper left zoom plot highlights the root mean square fluctuations between residue L200 and residue D230.

In order to highlight the large-scale and low frequency modes of cytochrome b, we performed the Principal Component Analysis (PCA). First, the Cα atoms of the cytochrome b were aligned, then the covariance matrix of the atomic fluctuations was calculated and diagonalized for each molecular system. The molecular trajectories were filtered along the first eigenvector obtaining the low frequency protein motion. Figure 5A shows an enlargement of the portion of cytochrome b under investigation and the average configurations representative of the phenomenon of interest obtained from the PCA analysis. The residues affected by these conformational changes are located near the Qi site, exposed to the mitochondrial matrix (Figure 5A) and their secondary structures are mainly random coils with a small portion of beta-sheet. During MD simulations, the highly unstructured portion of cytochrome b undergoes conformational changes by moving from a ‘closed’ to an ‘open’ state under the effect of thermal fluctuation (from black to white representation in Figure 5A). This mechanism is specific to the ligands present at the Qi site as highlighted by the mean RMSF (Figure 5B). The presence of Qi inhibitors favors the ‘closed’ state in a similar way to the apo form and the ubiquinone substrate (black representation in Figure 5A). In contrast, Qo inhibitors and the ubiquinol substrate favor a more frequent transition from closed to open state.

**Figure 5.**
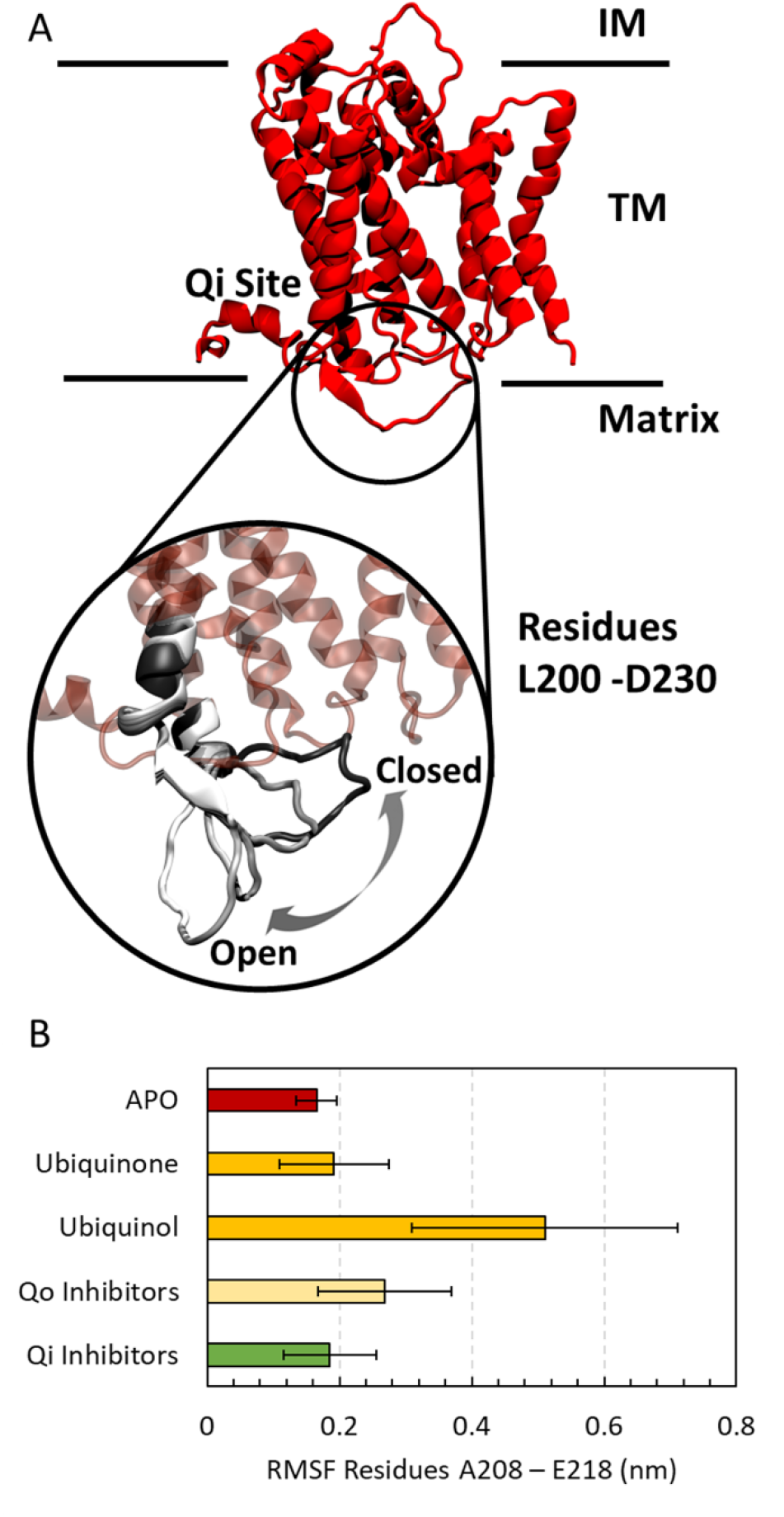
A) Heterodimeric cytochrome b structure representation. The three regions occupied by the receptor are the intermembrane space (IM), the transmembrane space (TM), and the matrix space. The residues from L200 to D230 near the Qi site are shown enlarged highlining the conformational changes by dynamically moving from a ‘closed’ (black cartoon) to an ‘open’ state (from gray to white cartoon). B) Root mean square fluctuations of the residues from A208 to E218.

The combined analysis of the RMSF and PCA suggests that the Qi inhibitors’ presence changes the rigidity in the proximity of the active site triggering the conformational changes of the cytochrome b of *T. cruzi*.

## Discussion

In the last decade, there has been renewed attention on Chagas’ disease caused by the *T. cruzi* due to the lack of well-validated molecular targets that limits the medical treatment(18,42–45). Cytochrome bc1 is a key component of the mitochondrial respiratory chain and a proven target for antimicrobial agents of medical and agricultural interest(11). Recent experimental and computational screening have identified the cytochrome bc1 and the production pathway of ubiquinone, through which cytochrome bc1 is highly dependent, as a prospective therapeutic target against T. cruzi(7,46,47). The cytochrome bc1 contains two catalytic sites (Qi and Qo), which are responsible for redox reactions that occur during the Q-cycle(16). Overall, a Q-cycle consumes two quinol molecules at the Qo site, creates one at the Qi site, and translocates four protons to the positive side of the mitochondrial membrane. Using a wide range of computational tools, we have investigated the mechanism of action of well-validated inhibitors for cytochrome bc1 by focusing on the Qi site. In detail, through homology modelling, molecular docking, and molecular dynamics simulations, we studied the impacts of four known inhibitors on cytochrome b conformational changes of *T. cruzi*. Two Qi-site inhibitors (antimycin A and GNF7686), two Qo-site inhibitors (myxothiazol and strobilurin A) were chosen and compared with two forms of substrate (ubiquinone and ubiquinol). Membrane proteins are the most accessible targets for therapies in receptor-based techniques, although structural information is limited. In this context, no crystal structure for the cytochrome bc1 of *T. cruzi* is available and homology modelling techniques have been employed. We have considered the cytochrome b heterodimer in order to focus on the Qi site. In particular, the homology model was built starting from the crystallography of *Gallus gallus*. In order to reproduce the physiological environment and dynamics of the receptor, the bL-heme and bH-heme molecules of the Qo and Qi catalytic sites were modelled, and the complex was placed in a neutral lipid bilayer model of POPC. The starting configuration of each ligand was predicted by molecular docking and the pose of the antimycin A ligand was used as a control for the docking protocol, obtaining RMSD values of less than 2 Å compared to the pose present in the crystallography with PDB ID 3H1I(25). The binding pocket and ligand reorganize during MD simulations, causing each ligand to interact with the receptor in a different way, affecting cytochrome b conformation. Overall, each ligand interacts mainly through hydrophobic interactions and forming hydrogen bonds with the residue Asp15 and Leu16. The Qo inhibitors coordinated via pi-staking iterations the residue Phe33 while antimycin A the residues His202 and Phe222 and gnf7686 the residue Phe194. Several structures containing quinone analogs showed the binding site to have several different configurations, and likely have different coordinating residues(48–50). The analysis of protein conformational changes revealed that the region between residue L200 and residue D230 has fluctuations specific to the type of ligand present at the Qi binding site. Overall, the investigated Qi inhibitors are capable of making this region more rigid than ligands related to Qo inhibitors and the same behavior is observed in the receptor apo form. The receptor’s apo form can in principle be considered as the receptor in an inactivated form. In light of this, we can hypothesize that the increased fluctuations of the highly unstructured portion under investigation may be due to the mechanism in which the cytochrome b works. Over time, computational studies have focused on understanding the mechanisms of drug action and resistance at the Qo site in order to shed light on the structure-activity relationship and improve drug design(21,22). The inhibitor stigmatellin binding at the Qo site near the ISP cluster [2Fe-2S] results in a decrease in the flexibility of the cytochrome bc1 complex(15,16). Furthermore, experimental studies highlighted the inhibitors’ action on the Qo and Qi sites changed the b-heme spectroscopic properties in cytochrome b(51–55) and coordination between the Qi and Qo sites influenced by Qi inhibitors was been hypothesised(56). We have schematized this movement as a fluctuation between a ‘closed’ and an ‘open’ state. The two substrates ubiquinone and ubiquinol exhibit opposite behavior. When ubiquinone occupies the Qi site, the average fluctuations are similar to those of the apo form, whereas ubiquinol causes large fluctuations up to 0.77 nm. In the last phase of the Q cycle, the ubiquinol substrate is produced by the reduction of ubiquinone at the Qi site, and in the last phase of the reduction process, two protons enter the intermembrane space. By considering these analyses, we propose that the fluctuation between a ‘closed’ and an ‘open’ state detected between residues L200 and D230 is assumed to be connected to and enhances the entry of protons into the intermembrane space. In detail, the inhibitors studied are specific for a particular binding site(13). The inhibitors act by competing with substrates and decreasing the biological activity of the cytochrome bc1 complex(15,16). In this context, the observed fluctuations might be a clear sign of the stability of the binding site associated with a given molecule, emphasizing inhibitors’ competitive effect against substrates and that Qo inhibitors are not suitable for inhibiting cytochrome bc1 activity by binding at the Qi site. In literature, atomistic MD simulations have hypothesized the role of cardiolipins as a source of protons during the redox reaction, suggesting that the overall mechanism could take into account several molecular factors(57). Experimental x-ray structure has emphasized the involvement of the highly conserved residue Arg218 in giving a proton during ubiquinone reduction and then retrieving it from the matrix(48). A similar hypothesis was made about a possible pathway of proton conduction out of the Qo site, that through a water channel in the cytochrome bc1 involved the bL-heme and connected the lobe near the bL-heme to the solvent(21). To summarize, the mechanism of action of inhibitors and receptor function is quite complex and involves numerous variables. Future studies, especially those aimed at identifying new inhibitors for the Qi site of cytochrome b, will be able to assess the validity of this model for developing more effective drugs.

## Data and software availability

Specific open access software from third parties was used: GROMACS 2020.4 (https://manual.gromacs.org/documentation/2020.4/download.html), VMD1.9 (http://www.ks.uiuc.edu/Research/vmd/), MODELLER (https://salilab.org/modeller/), ProLIF (https://prolif.readthedocs.io/en/latest/index.html), PROCHECK (https://www.ebi.ac.uk/thornton-srv/software/PROCHECK/), T-Coffee algorithm (https://www.ebi.ac.uk/Tools/msa/tcoffee/), PLANTS and SPORES (http://www.tcd.uni-konstanz.de/plants_download/). Parameter files are publicly available from http://mackerell.umaryland.edu/charmm_ff.shtml. Molecular structures used are available from https://www.rcsb.org/. Trypanosoma cruzi strain Silvio kinetoplast maxicircle sequence is available from https://www.ncbi.nlm.nih.gov/nuccore/FJ203996.1. Output files are available from the corresponding author upon request.

## Author Information

### Author Contributions

S.M and G.G conceived the research. S.M did the homology modelling, molecular docking, and the molecular dynamics simulations. S.M, G.G, L.S and AD analyzed and rationalized the data. All authors wrote the paper and critically commented to the manuscript for important intellectual content. All authors read and approved the final manuscript.

## Acknowledgments

This work was supported by a grant from the Swiss National Supercomputing Centre (CSCS).

## Abbreviations

MD: molecular dynamics
ETC: electron transport chain
PCA: Principal Component Analysis
VMD: Visual Molecular Dynamics
bL-heme: low potential b heme
bH-heme: high potential b heme
RMSF: root mean square fluctuations.

## Supporting information

**Table S1**: **Molecular systems investigated**. (PDF)

**Figure S1**: **Sequence alignment**. Sequence alignment of cytochrome b of T. cruzi, and G. gallus. (TIFF)

**Figure S2**: **Homology model and template model**. A) Cartoon representation of the X-ray structure of Cytochrome b of Gallus gallus (PDBID 1BBC). The binding Qi site is shown with cyan surface on the ubiquinone natural compound. B) Cartoon representation of superposition between the avian cytochrome b template structure (red) and the T. cruzi cytochrome b homology model (green). (TIFF)

**Figure S3**: **Ramachandran plot**. Psi-phi angle in degree of the cytochrome b homology model built. (TIFF)

**Figure S4**: **Representative complex model**. A) Representative complex model built and B) an enlargement of the ligand and heme groups. The cytochrome b is shown in orange, the POPC lipid bilayer in violet, the heme groups in cyan and the ligand in the Qi site in green. (TIFF)

**Figure S5**: **Molecular docking validation**. Molecular docking pose validation of the antimycin A in the Qi site. (TIFF)

